# Estimating the age of poorly dated fossil specimens and deposits using a total-evidence approach and the fossilized birth-death process

**DOI:** 10.1101/2021.04.12.439507

**Authors:** Joëlle Barido-Sottani, Dagmara Żyła, Tracy A. Heath

## Abstract

Bayesian total-evidence approaches under the fossilized birth-death model enable biologists to combine fossil and extant data while accounting for uncertainty in the ages of fossil specimens, in an integrative phylogenetic analysis. Fossil age uncertainty is a key feature of the fossil record as many empirical datasets may contain a mix of precisely dated and poorly dated fossil specimens or deposits. In this study, we explore whether reliable age estimates for fossil specimens can be obtained from Bayesian total-evidence phylogenetic analyses under the fossilized birth-death model. Through simulations based on the example of the Baltic amber deposit, we show that estimates of fossil ages obtained through such an analysis are accurate, particularly when the proportion of poorly dated specimens remains low and the majority of fossil specimens have precise dates. We confirm our results using an empirical dataset of living and fossil penguins by artificially increasing the age uncertainty around some fossil specimens and showing that the resulting age estimates overlap with the recorded age ranges. Our results are applicable to many empirical datasets where classical methods of establishing fossil ages have failed, such as the Baltic amber and the Gobi Desert deposits.

## 1 Introduction

Recent progress in statistical methods has enabled biologists to estimate the timing of speciation events in phylogenies comprising both living and fossil taxa. These advances include likelihood-based models for discrete morphological data—variants of the Mk model (Lewis, 2001)—that describe the substitution process for discrete character data, and thus allow for statistical inference of phylogenetic relationships from morphological matrices. When combined with models characterizing the distribution of substitution rates among branches (such as “relaxed clock” models like those described by Thorne et al., 1998; Drummond et al., 2006; Lepage et al., 2007, and many others), these advances led to the introduction of new Bayesian approaches for jointly estimating phylogenetic relationships and divergence times of datasets containing extant taxa and dated fossil specimens. Early applications of these Bayesian “total-evidence” dating analyses (Pyron, 2011; Ronquist et al., 2012a) did not adequately model the speciation-extinction-sampling process underlying the generation of a dated phylogenetic tree with sampled fossil and extant taxa (Pett and Heath, 2020). However, the serially sampled birth-death process introduced by Stadler (2010) was later integrated into Bayesian approaches for inferring time-calibrated phylogenies using more realistic models of diversification and sampling (Heath et al., 2014; Gavryushkina et al., 2014). This model is referred to as the fossilized birth-death (FBD) process when applied to datasets including information from the fossil record (Heath et al., 2014).

The FBD process describes the generation of a dated phylogenetic tree of sampled extant and fossil lineages, with parameters explicitly controlling for the extant sampling probability and the rates of speciation, extinction, and fossil recovery. This model can be combined with the morphological and clock models described above in a Bayesian statistical framework. Moreover, this integrative Bayesian framework allows researchers to combine paleontological information into phylogenetic analyses of living species, thus providing insights into the timing and rate of diversification in the tree of life. Importantly, total-evidence methods using the FBD model allow researchers to include a greater amount of the data observed from the fossil record, which, in turn, improves our understanding of macroevolutionary processes. Bayesian total-evidence methods and associated models are implemented in statistical tools like RevBayes (Höhna et al., 2016), BEAST2 (Bouckaert et al., 2014, 2019), and MrBayes (Ronquist et al., 2012b). With access to statistical software for more holistically integrating paleontological and neontological data, biologists have greatly improved our understanding of the evolutionary dynamics of various clades including monocots (Eguchi and Tamura, 2016), beetles (Gustafson et al., 2017), sponges (Schuster et al., 2018), vipers (Šmíd and Tolley, 2019), and termites (Jouault et al., 2021).

The fossil record is essential for calibrating species trees to time (*i*.*e*., years or millions of years), as molecular sequences from extant species are informative about the relative age of species but do not typically provide information about the absolute age (Pett and Heath, 2020). There are two main methods of determining a fossil’s age, namely relative dating and absolute dating. Relative fossil dating determines a specimen’s approximate age by comparing it to similar rocks and fossils with known ages. A fossil’s absolute date is obtained by applying radiometric dating to measure the decay of isotopes, either within the fossil or, more often, the rocks associated with it (Gradstein et al., 2012; Peppe and Deino, 2013). Accurate dates for fossil specimens and deposits are critical not only for understanding the timing of speciation events in the tree of life, but these dates also provide crucial data for answering questions in evolutionary biology, paleoecology, biogeography, and paleoclimatology. However, there are deposits and key specimens where traditional dating methods have failed and their ages remain uncertain. Uncertain dates for fossil specimens and formations, in turn, limit the scientific value of these observations.

One of the most famous examples of such a deposit is Baltic amber, a remarkable source of terrestrial invertebrate fossils (mostly insects) from the Eocene. There are several hypotheses concerning its age (for a summary see Bogri et al., 2018) and it is generally dated as Eocene, with a wide age range between 55 and 34 Ma. Difficulties in the age determination are due to the repeated re-deposition of the amber, the broad range of the ancient forest, and its probable existence for several million years. Another example where the age uncertainty hampers biological and geological studies is the Cretaceous terrestrial sediments in the Gobi Desert of Mongolia, a site renowned for remarkably well preserved vertebrate fauna, including dinosaurs. Unfortunately, a definitive age cannot be directly determined due to the lack of discrete key beds, like zircon-bearing tuffs (Kurumada et al., 2020). In some cases, even if the age range of a formation can be determined, other factors might hinder the assessment of a fossil’s age. One example of such a deposit is the Daohugou Formation (164-159 Ma), which is well known for exceptionally complete fossils, including a diverse and rich record of invertebrates and plants, but also many vertebrates preserved with traces of soft tissues (Wang et al., 2005). However, due to the complicated stratigraphy of the formation, where several fossiliferous layers mix and overlap (Li et al., 2021), it is often difficult to assess a fossil’s precise age without knowing the exact layer from which it was sampled.

Without sufficient direct evidence for dating critical deposits and specimens, scientists must rely on approaches that harness the information in indirect evidence. Bayesian total-evidence approaches make it possible to directly integrate the age uncertainty around historic samples into Bayesian analyses (Shapiro et al., 2011) and previous work has shown that adequately representing this uncertainty is critical to obtaining accurate phylogenies and divergence times estimates (Barido-Sottani et al., 2019a, 2020b). However, most phylogenetic divergence-time analyses typically treat fossil ages as nuisance parameters and the uncertainty associated with those observations is simply a source of error. Nevertheless, the ages of heterochronous specimens may be particularly interesting for some types of phylogenetic studies. Shapiro et al. (2011) note that datasets of infectious diseases or those that include ancient DNA sequences may have samples with unknown ages, and robust estimates of these undated samples can help shed light on the dynamics of viral epidemics or the ecological contexts of sub-fossils used in ancient DNA research. Their simulations and empirical validations show that phylogenetic analyses of datasets including a single undated sample can yield accurate estimates of the unknown sampling time (Shapiro et al., 2011). More recently, Drummond and Stadler (2016) extended this study to consider much older time-scales and total-evidence analyses of fossil and extant species under the fossilized birth-death model. Their study focused on analyses of fossil-rich empirical datasets and demonstrated that the age estimated for a single fossil specimen with an unknown date is accurate when using this integrative Bayesian approach (Drummond and Stadler, 2016). While these previous studies indicate that combining data from extant and fossil taxa can lead to accurate age estimates for poorly dated fossils, they did not consider the patterns of age uncertainty frequently associated with the fossil record. Paleontological datasets can often include collections of fossils all sampled from the same poorly dated formation or multiple fossils with incomplete or disputed provenance, making it difficult to assign an accurate date.

In this study, we investigate the performance of Bayesian phylogenetic approaches using the FBD model, applied to datasets that include multiple fossils from poorly dated formations. We use simulations to evaluate the accuracy and robustness of the age estimates for fossils belonging to the uncertain formation, and explore whether the presence of poorly dated fossils affects the estimates of the tree topology and the ages of the other, well dated fossils. We use a recently published dataset of extant and fossil penguins (order Sphenisciformes) from Thomas et al. (2020) to validate this approach on empirical data. Fossil penguin specimens have relatively precise dates, allowing us to compare the age estimates obtained when artificially increasing the age uncertainty around some selected fossils to ages observed and recorded from the fossil record.

## 2 Methods

### 2.1 Simulated data and analyses

We evaluated the accuracy and precision of fossil specimen age estimates using simulated datasets. We calibrated the model and parameters used for simulation based on an empirical dataset of the subfamily Paederinae of rove beetles (Staphylinidae, Coleoptera). This subfamily has a strikingly rich fossil record in Cenozoic deposits, including several fossil specimens from one of the best known poorly dated insect deposits, Baltic amber (DŻ, personal observations), making rove beetles well suited for providing realistic values for our simulations.

### 2.2 Simulated phylogenies and taxon sampling

Trees were simulated under a birth-death process using the R package TreeSim (Stadler, 2011), starting from one lineage at the origin time of 120 Myr, with the speciation rate set to *λ* = 0.05/Myr and the extinction rate to *µ* = 0.02/Myr. Speciation and extinction rates were selected based on estimates for the Staphylininae subfamily of rove beetles, from Brunke et al. (2017). For each simulation condition, 100 replicates were simulated. The extant sampling probability was set to *ρ* = 0.5. In order to keep the trees computationally manageable, we based the number of tips based on the number of genera currently classified into Paederinae (A. Newton, unpublished database). Thus we rejected trees which had less than 20 or more than 30 extant samples.

In our setup, we assume that the unknown deposit is likely tied to a geographical or ecological factor affecting the corresponding lineages. Thus to sample fossils, we first assigned all lineages present in the complete tree falling within the 30 and 50 Myr interval to a binary character, using a continuous rate transition process where all lineages started in state 1 at age=50 and transitioned from state 1 to 2 with rate *q*_1,2_ and back with rate *q*_2,1_. All lineages occurring outside of the 30 *−* 50 interval were assigned to state 1. We then sampled fossils using the R package FossilSim (Barido-Sottani et al., 2019b), following a Poisson process with piece-wise constant rates *ψ*_*int*_ between 30 and 50, and *ψ*_*bg*_ outside of this interval. Fossil samples in state 1, designated as “precise-date” fossils, were considered to be individual samples, while fossil samples in state 2, designated as “imprecise-date” fossils, were assigned to all occur within the same poorly dated deposit. Transition and fossilization rates were calibrated to obtain specific proportions (0.1, 0.3 or 0.5) of imprecise-date samples among all fossils. The detailed values used are shown in Table 1. Simulations were rejected if the resulting proportion was more than 10% different from the target proportion, or if the total number of fossil samples was not between 45 and 55. Note that in order to obtain the target proportions, a higher sampling rate had to be used during the interval of sampling imprecise-date fossils compared to the rest of the timeline.

**Table 1:** Parameter values used to simulate the fossil sampling process.

| Target proportion of<br>imprecise-date fossils | $q_{1,2}$ | $q_{2,1}$ | $\psi_{bg}$ | $\psi_{int}$ |
| --- | --- | --- | --- | --- |
| 0.1 | 0.6 | 0.7 | 0.03 | 0.04 |
| 0.3 | 0.8 | 0.5 | 0.02 | 0.08 |
| 0.5 | 1.0 | 0.4 | 0.01 | 0.15 |

An example of a complete simulated tree with fossil samples is shown in Figure S4. To simulate fossil age uncertainty, all fossil samples were assigned a range of possible ages, depending on their state. Imprecise-date fossils were all assigned the same age range of 30 to 50 Myr, and precise-date fossils were assigned a range of fixed length 0.1, 0.2 or 0.3 times the true age of the fossil. The minimum age of each range was sampled uniformly so that the true age of the fossil always lied within its corresponding range.

#### 2.1.2 Molecular sequence alignment and morphological character matrix simulation

Molecular sequences were simulated for the extant samples using seq-gen (Rambaut and Grassly, 1997) via the R package phyclust (Chen, 2011). We simulated sequences comprising 4,500 nucleotides under an HKY+Γ model with five rate categories and a gamma shape value of *α* = 0.35. As the inference of the phylogenetic tree from molecular data was not the focus of this study, we used a simple strict molecular clock, with a clock rate set to 0.05 substitutions/Myr, based on estimates of the clock rate from Brunke et al. (2017).

Morphological alignments were simulated for both extant and extinct samples using the R package geiger (Pennell et al., 2014). We simulated matrices of 120 characters under an Mk model (Lewis, 2001) with five rate categories, selecting only varying characters. The number of states in the simulated matrices varied such that 70% of simulated characters were binary, 20% ternary, and 10% quaternary. The morphological clock rate was set to 0.1 substitutions/Myr, following an estimate for Chrysomelidae and Cerambycidae from Farrell and Sequeira (2004). We assigned a random proportion of 5% of the simulated morphological characters as “soft” characters, which were only represented in extant taxa and were assigned the unknown character “?” for all fossil samples, thus emulating biased character preservation.

#### 2.1.3 Bayesian inference

For each simulated dataset, we performed a Bayesian total-evidence analysis in RevBayes (Höhna et al., 2016) under a constant-rate FBD tree prior. The constant-rate FBD model is used in most empirical studies, as time-dependent variation in rates is often difficult to know *a priori*. Priors for the speciation, extinction, and fossilization rates were set to Exponential(10). The ages of the fossils were sampled along with the other parameters, with a prior set as uniform over their simulated range, as described in Drummond and Stadler (2016). The extant sampling proportion was fixed to the true value, *ρ* = 0.5. Moves were set in accordance with guidance from the RevBayes FBD tutorial (Barido-Sottani et al., 2020a, also see: https://revbayes.github.io/tutorials/fbd/fbd_specimen.html). The substitution and clock models were set to the simulation models. The parameters of these models were estimated, using priors and moves also set following the RevBayes FBD tutorial. The full Rev scripts used for inference are available in the Supplementary Materials. Analyses in RevBayes were run for up to 150,000,000 generations, and two independent chains were run in for each replicate. Samples from each run were assessed in Tracer (Rambaut et al., 2018). We considered that the Markov chain had reached stationarity and converged on the target distribution if the effective sample size (ESS) of the posterior had reached a value > 200 and if both chains had median posteriors which differed by no more than 10%. We did not assess the convergence of the tree topology. Some simulation replicates (0 to 12 depending on the dataset, out of 100) failed to converge and were discarded from the final results.

#### 2.1.4 Assessing results

We assessed the accuracy of the fossil age estimates by measuring the relative error of the posterior estimates, defined as the absolute difference between the true value and the estimated value, divided by the true value. We also calculated the coverage, *i*.*e*., the proportion of analyses in which the true parameter value was included in the 95% highest posterior density (HPD) interval. These measures were averaged separately over all imprecise-date and precise-date fossils.

To assess the accuracy of inferred topologies we calculated the mean normalized Robinson-Foulds (RF) distance (Robinson and Foulds, 1981) between the true simulated trees, including the fossil samples, and the tree samples from the posterior distribution. The RF distance only depends on the topology of the trees. The normalized RF distance between two trees with *n* tips is computed by dividing the RF distance between these trees by the maximum possible RF distance between two trees with *n* tips, thus scaling the distances between 0 and 1. Finally, we assessed the accuracy of the positioning of fossils on the inferred tree topologies by calculating the proportion of posterior samples in which a given fossil was placed in the correct extant clade.

### 2.2 Validation using empirical data

We used a recently published study of penguins by Thomas et al. (2020) to demonstrate how Bayesian phylogenetic analyses can improve the precision of poorly dated fossil specimens using an empirical dataset. This dataset is a useful “ground truth” for fossil age estimation because the extant diversity of penguins is completely sampled, which minimizes the effect of potential sampling biases in the analysis. Moreover, the majority of fossils in this dataset are precisely dated (age ranges of 1.5 to 10 My) and the penguin fossil record is generally considered reliable.

We used the molecular and morphological data matrices of living and fossil Sphenisciformes published in Thomas et al. (2020), which include recently published sequences from Cole et al. (2019) and extend the morphological matrix by Degrange et al. (2018). The molecular sequence alignment contains mitochondrial genome sequences of 15,755 nucleotides for 24 extant taxa, and the morphological matrix is composed of 274 characters for 66 extant and fossil species. We focused our study on the estimated ages of fossil taxa while marginalizing over the tree topology (for the tree topology see figure 2 in Thomas et al., 2020).

The observed age ranges for all fossil species were obtained from Thomas et al. (2020). We imposed a poorly dated fossil deposit on this dataset by assigning an identical large age range to selected fossil species. The observed age range of the fossils was always fully included in the assigned age range. Unlike the simulated dataset, we did not use the age range of the Baltic amber deposit. Instead, we selected three age intervals which covered the age ranges of approximately the same number of species, but were of different length. The first interval, denoted as “small”, ranged from 30.3 Ma to 46.8 Ma and contained 14 fossil species. The second interval, denoted as “large”, ranged from 14.6 Ma to 44.6 Ma and contained 15 fossil species. We also tested an extension of the first interval, which ranged from 25.2 Ma to 61.5 Ma and contained 22 fossils species. For each interval, two conditions were tested: (1) a random subsample of 50% of the species were assigned the full interval as age range, while the other species were assigned their observed ranges; and (2) all fossil species in the interval were assigned the full interval as their age range. In contrast to the simulation setup, the assignment of fossils to the unknown deposit was not tied to a phylogenetic character. The full prior age ranges set for each fossil and each configuration is shown in Figures S8-S10.

#### 2.2.1 Bayesian inference

We performed the phylogenetic analyses in RevBayes (Höhna et al., 2016). With the exception of the age ranges, which were modified as described in above, all models and priors were identical to the analysis in Thomas et al. (2020), which also used the RevBayes FBD tutorial as a guide (Barido-Sottani et al., 2020a, also see: https://revbayes.github.io/tutorials/fbd/fbd_specimen.html). All fossil ages were assigned a uniform prior distribution over their age range. Priors for the speciation, extinction, and fossilization rates were set to Exponential(10). The molecular alignment used a *GTR* + Γ substitution model with 4 rate categories, in combination with an uncorrelated exponential clock model with a prior of Exponential(10) on the mean clock rate. The morphological alignment used an Mk substitution model (Lewis, 2001) with 4 rate categories, in combination with a strict clock model with a prior of Exponential(1) on the clock rate. The inference was run for 137,000,000 iterations. Convergence was assessed in Tracer (Rambaut et al., 2018) using the criteria described above for the simulation analyses.

## 3 Results

### 3.1 Simulated datasets

Results from our analyses of the simulated datasets are shown in Figures 1 and 2. As expected, the relative error on age estimates is much higher for imprecise-date fossils than for precise-date fossils. The proportion of imprecise-date fossils has a strong impact on the accuracy of fossil age estimates, with the mean coverage for the estimated age of imprecise-date fossils ranging from *≈* 65% when 10% of the fossils are poorly dated, to only *≈* 30% when 50% of the fossils are poorly dated. However, the absolute relative error remains quite low for imprecise-date fossils even in the worst case scenario, indicating that the inference is able to recover approximate age estimates for fossils from poorly dated deposits, despite the decreased coverage. The width of the age range associated with precise-date fossils, which corresponds to the magnitude of the age uncertainty on those fossils, has a strong impact on the accuracy of the age estimates for well dated fossils, but little effect on the estimates for imprecise-date fossils. This holds true even in the datasets where the relative age range for precise-date fossils is 30%, and the oldest precise-date fossils are associated with more age uncertainty than imprecise-date fossils. One likely reason for this is that older fossils are relatively rare in our simulated datasets, for instance only *≈* 15% of the total number of fossils are older than 60My.

**Figure 1.**
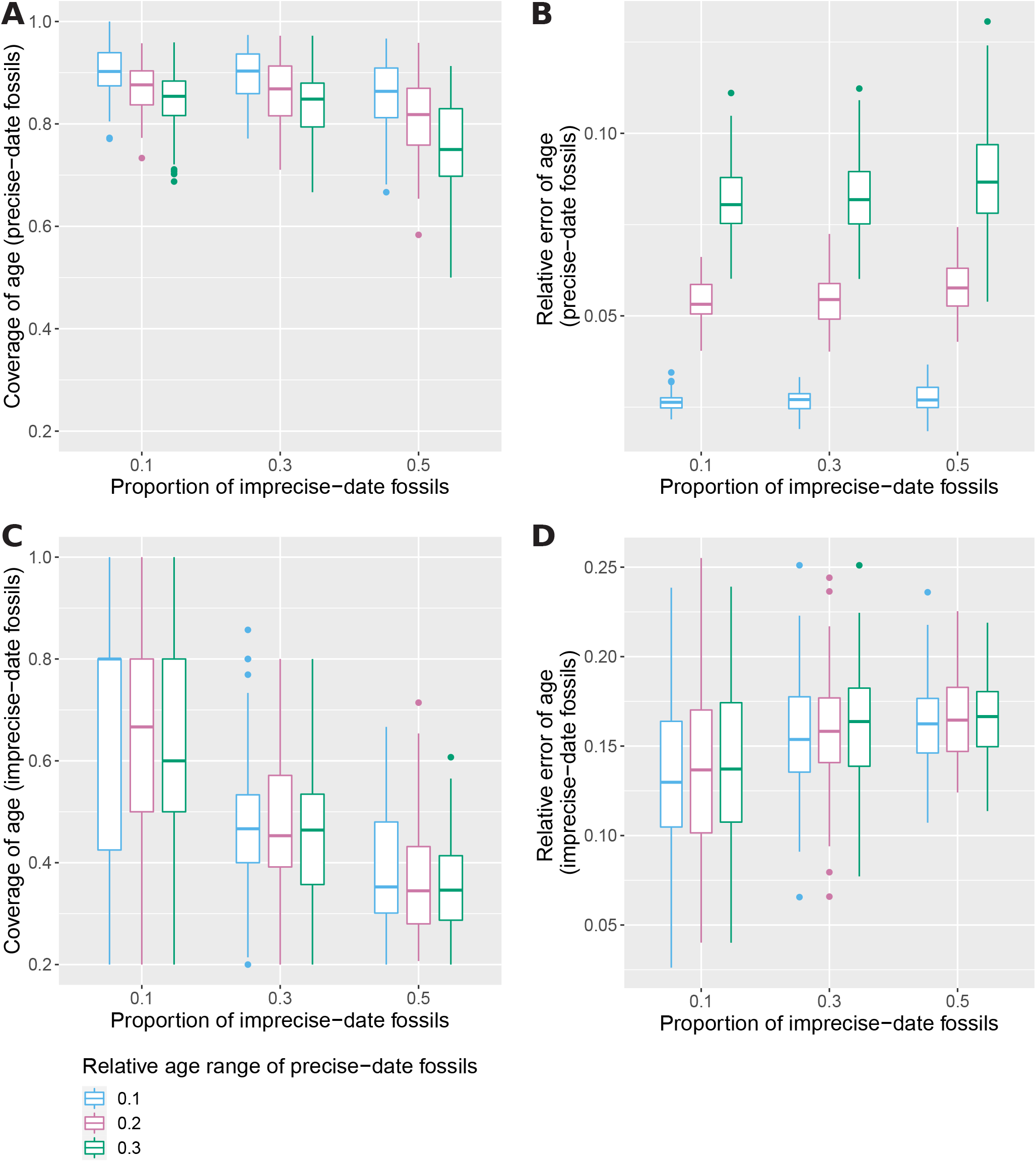
Relative error of the median age estimate (B,D) and 95% HPD coverage (A,C) of precise-date fossils (A,B) and imprecise-date fossils (C,D) for different proportions of imprecise-date fossils, and different widths of the age range of precise-date fossils. Measures are averaged over all fossils for each replicate. The average and standard deviation across all replicates is shown.

**Figure 2.**
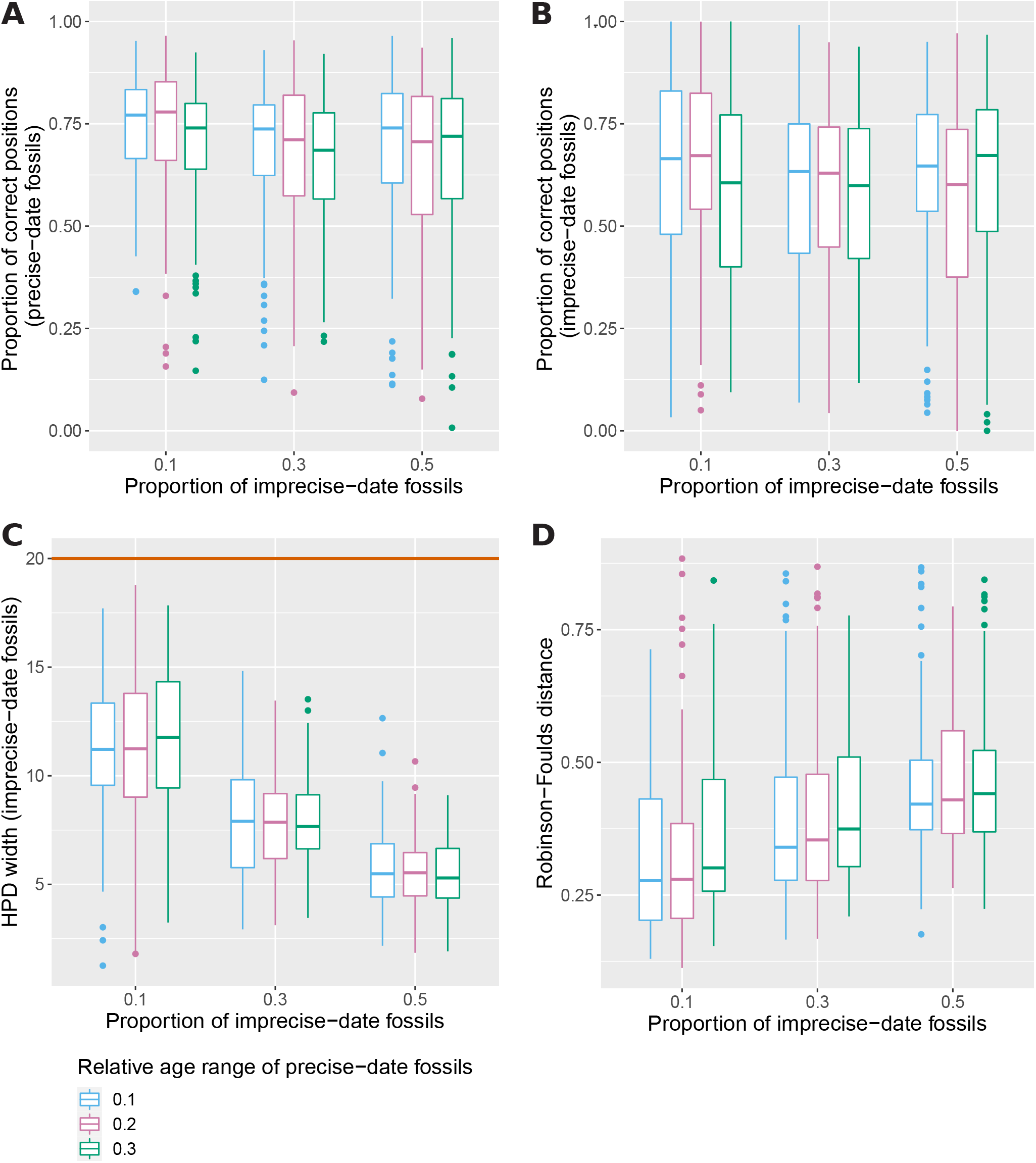
Proportion of posterior samples with correctly placed fossils, averaged across all precise-date fossils (A) or all imprecise-date fossils (B), width of the 95% HPD interval averaged across all imprecise-date fossils (C) and mean normalized RF distance between estimated trees and simulated tree (D), for different proportions of imprecise-date fossils and different widths of the age range of precise-date fossils. The average and standard deviation across all replicates is shown. The brown line in C shows the size of the age range set as the prior for all imprecise-date fossils (*i*.*e*., 20Myr).

The widths of the 95% HPD intervals for imprecise-date fossils are smaller than the time interval of the prior age range in all tested conditions, showing that the age estimates of imprecise-date fossils are not driven only by this prior (Fig. 2C). Interestingly, the HPD interval widths decrease with higher proportions of imprecise-date fossils, while the estimates show decreased accuracy and coverage in this situation. This is contrary to the expected pattern, which would be that interval widths increase with larger amounts of uncertainty in the data, but that coverage levels remain similar. One likely explanation is that our simulations used a piece-wise constant sampling rate, in violation of the inference model which assumes that all FBD rates are constant across time and lineages. In addition, the discrepancy between the low and high fossil sampling rate increased with the proportion of imprecise-date fossils. It is also likely that the impact of model violation on the estimates is stronger in datasets with lower amounts of data. The combination of these two factors leads the datasets with high proportions of fossils with imprecise dates to exhibit narrower than expected HPD intervals and decreased coverage.

The topology inference follows a similar pattern, as the proportion of correct fossil positions decreases and the average RF distance increases with increasing age uncertainty or higher proportions of poorly dated fossils (Fig. 2D). The positions of precise-date fossils are more accurate than the positions of imprecise-date fossils, which confirms that fossil ages are an important source of information for both topology and branch times in total-evidence analyses. In all tested conditions, the average proportion of correct fossil positions is *>*50% for both precise- and imprecise-date fossils.

### 3.2 Empirical dataset

Figure 3 shows the results of the analysis on the penguin datasets, using either the small or the large interval as deposit. When only 50% of the fossils in the interval are given imprecise dates (Fig. 3A and C), there is a large overlap between the estimated posterior distributions of fossil ages and the empirical intervals. As expected, the overlap decreases when all fossils in the interval have imprecise dates and less information is available in the dataset (Fig. 3B and D). In this case, some posterior distributions diverge completely from the recorded interval (Fig. 3D, *Paraptenodytes antarcticus*), or appear to be driven mostly by the prior (Fig. 3B, *Delphinornis arctowskii*).

**Figure 3.**
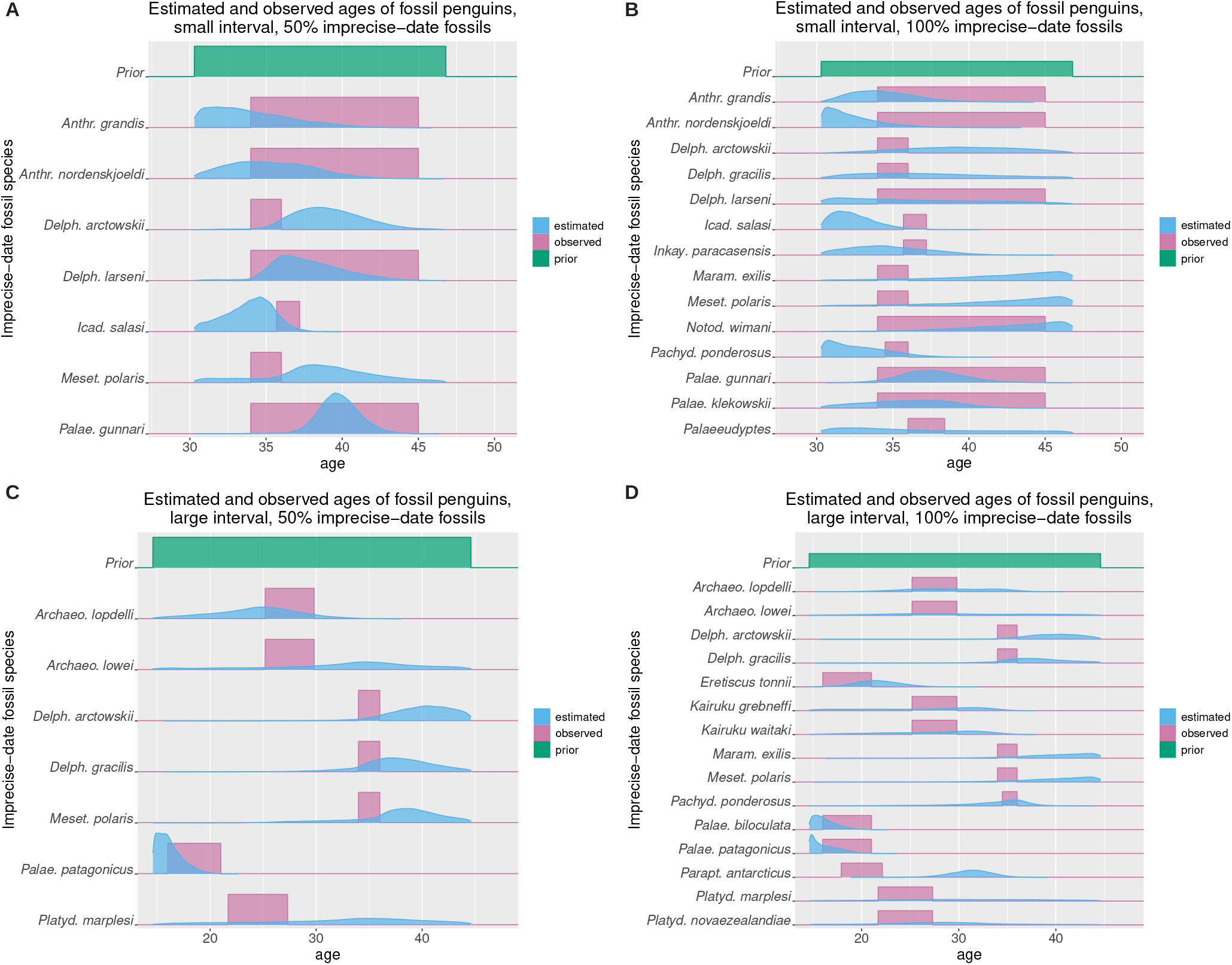
Comparison of observed (pink) and estimated (blue) penguin ages for the small (A,B) and large (C,D) intervals, with a proportion of 0.5 (A,C) or 1 (B,D) of imprecise-date fossils. The observed age range is shown as a uniform distribution, while the estimated age is the inferred posterior distribution. The uniform distribution used as prior for the imprecise-date fossils is shown in green on each panel.

The results concerning the extended version of the first interval are shown in the supplementary materials, and show similar patterns to the large interval. Overall, these results confirm that the age of the well dated fossils, in combination with the tree, allows us to estimate the age of poorly dated fossils, and that the presence and number of these well dated fossils plays a key role in the accuracy of the resulting estimates.

## 4 Discussion

While phylogenetic analyses using the FBD model have largely focused on inferring phylogenetic trees and dating species divergences, our study shows that these methods can harness indirect information in an integrative and hierarchical model to improve date estimates for fossil specimens themselves. This is also the case when the dataset includes a collection of poorly dated fossils that all come from the same formation. This showcases one of the strengths of the FBD process as a complete model integrating both diversification and fossil recovery processes.

Our study examines the accuracy of age estimates for a combination of poorly dated fossils from the same deposit and more credible fossils from well dated deposits. We show that when these fossil taxa are integrated with extant species in a joint analysis of discrete morphological characters (fossil and extant) and molecular sequences (extant only), it is possible to infer the ages of fossil samples from a deposit with a large age uncertainty. As expected, the accuracy of the fossil age inference is strongly impacted by the amount of uncertainty and missing information present in the analysis, which is represented in our study by the relative proportion of fossils with uncertain dates versus those with precise dates, as well as the magnitude of the age uncertainty associated with well dated fossils. Finally, we also demonstrate that the extant topology and the overall age of the phylogeny are well estimated in all datasets, which shows that FBD total evidence analyses can provide reliable estimates despite including fossils with large amounts of age uncertainty.

It is important to note that our simulations represent an idealized scenario, chosen to reduce model complexity and the noise of the parameters under examination, and to focus specifically on fossil age estimates. In particular, we used a strict clock for both the molecular and the morphological alignments, which is likely to be unrealistic for large empirical datasets. As shown in the supplementary materials, using a relaxed clock for the molecular alignment did not significantly affect our results, but also led to convergence issues, particularly in combination with high proportions of imprecise-date fossils. Using a relaxed clock for the morphological alignment also led to reduced sampling efficiency and poor convergence. In general, we expect that increasing the complexity of the model can induce long mixing times for the Markov chain Monte Carlo (MCMC) sampler and in some cases lead to non-convergence. One potential way to reduce complexity would be to assign the same age to all imprecise-date fossils, rather than estimating all fossil ages independently as we did. However, this assumes that all fossils from the deposit were sampled at very similar dates, rather than the deposit being the product of a continuous fossilization process over an extended period of time. In practice, it may be difficult to distinguish between these two hypotheses *a priori*. The Baltic amber deposit, which we use as the basis for our simulation parameters, is particularly challenging in this respect. This deposit is an umbrella term for various secondary amber deposits found around the Baltic Sea. It remains unclear whether amber found in the different regions originated in a single or in multiple areas. The north European Eocene forest covered a substantial area of many square kilometres and amber forests in the Baltic region could have persisted for several million years up to the end of the Eocene (for a summary see Bogri et al., 2018).

Poor mixing and convergence issues are particularly problematic when a complex, parameter-rich model is applied to a dataset with large amounts of missing data. As a result, we expect that this approach may perform less well than our simulated results on more complex empirical datasets, and in some cases may not converge at all without careful attention to the MCMC proposal algorithms. We believe that this is inevitable due to the challenges of working with missing data. Other ways to reduce uncertainty and complexity may be used, such as topological constraints which use taxonomic information to place fossil samples in particular clades, instead of relying purely on the morphological data and fossil ages to inform the inference of the tree topology. These constraints are particularly helpful in datasets where available morphological matrices are small (*<* 50 characters), since previous work has shown that small morphological matrices lead to high levels of inaccuracy in topology estimates (Barido-Sottani et al., 2020b). In addition, one advantage of using a Bayesian approach is that estimates will accurately represent the amount of uncertainty present in the dataset under a given model, including cases where the amount of uncertainty is too large to draw exploitable conclusions. However, this is only true if the inference model matches with the true evolutionary process, or in our case, with the simulation model.

One likely contributor to the decrease in accuracy when the proportion of imprecise-date fossils increases is that our inferences assumed uniform fossil sampling rates throughout the tree, an assumption which was increasingly violated when the proportion of fossils coming from the same deposit increased. The assumption of uniform sampling is very uncommon in existing empirical analyses, however our results show that using this assumption when a large proportion of fossils come from the same deposit can lead to biases in the inference. Therefore, we advise empirical studies to pay attention to the time and spatial distribution of the included fossils, and to use the skyline FBD model (Stadler et al., 2013; Zhang et al., 2016) if time-varying rates are a likely factor.

Understanding the performance of statistical phylogenetic methods under realistic conditions is especially critical for methods applied to paleontological data. The structure and complexity of the geologic record (Holland, 2016) as well as the challenges associated with collecting and curating fossils that may lead to uncertainty in a specimen’s age, collection locale, or identification are all common realities faced by researchers working with fossils. Thus, new phylogenetic models that account for the way that taxa are sampled (*e*.*g*., Höhna et al., 2011) or how fossil data are influenced by the structure of the rock record (e.g., Stadler et al., 2018) will be important for improving our understanding of the geological and ecological context of lineage diversification through time.

In conclusion, we show that total-evidence phylogenetic analyses under a fossilized birth-death process can improve the precision of age estimates for fossils sampled from poorly dated geologic formations when combined with character data and other information from extant taxa and other well dated fossil species. This approach may be useful for empirical datasets where the majority of fossils are precisely dated, but some specimens are sampled from a deposit with uncertain dates, *e*.*g*., the Baltic amber deposit for insects (such as the rove beetles used as a model in our study) or the Gobi Desert deposit for dinosaurs. Such analyses are easily extended to include other processes present in empirical data, such as diversified sampling of extant taxa (Höhna et al., 2011), which accounts for taxonomy-guided sampling strategies where only a single representative per genus or family is included in a dataset. However, because the accuracy of parameter estimates may be reduced when such complex models are used in analyses of highly incomplete datasets, researchers applying these methods to estimate fossil ages are encouraged to consider ways where they can minimize uncertainty and increase sampled data. Importantly, for some taxonomic groups, this may require more support and time for efforts to collect, curate, and analyze paleontological data.

## Supporting information

Supplementary methods and results

## Acknowledgements

We thank R. Warnock and J. Satler for helpful comments on this manuscript. We are grateful to A. Brunke for sharing his files from the Staphylininae dating analysis. JBS and TAH were supported by funds from the National Science Foundation (USA), grants DBI-1759909 and DEB-1556615. This project has received funding from the European Union’s Horizon 2020 Research and Innovation Programme under the Marie Sklodowska-Curie grant agreement No. 797823 (postdoctoral fellowship of DŻ) and No. 101022928 (postdoctoral fellowship of JBS). All analyses were conducted using the Nova cluster, part of the Iowa State University High Performance Computing resources.

## Data availability

The R scripts used for simulation, post-processing and plotting, the Rev scripts used for running the inference, as well as the simulated and empirical data files are available as a Zenodo repository, DOI: 10.5281/zenodo.6902473.

## References

Barido-Sottani, J., G. Aguirre-Ferńandez, M. J. Hopkins, T. Stadler, and R. Warnock. 2019a. Ignoring stratigraphic age uncertainty leads to erroneous estimates of species divergence times under the fossilized birth-death process. Proceedings of the Royal Society B: Biological Sciences 286:20190685.

Barido-Sottani, J., J. A. Justison, A. M. Wright, R. C. Warnock, W. Pett, and T. A. Heath. 2020a. Estimating a time-calibrated phylogeny of fossil and extant taxa using RevBayes. Pages 5.2:1–5.2:23 in Phylogenetics in the Genomic Era (C. Scornavacca, F. Delsuc, and N. Galtier, eds.). No commercial publisher | Authors open access book.

Barido-Sottani, J., W. Pett, J. E. O’Reilly, and R. C. Warnock. 2019b. FossilSim: an R package for simulating fossil occurrence data under mechanistic models of preservation and recovery. Methods in Ecology and Evolution 10:835–840.

Barido-Sottani, J., N. M. van Tiel, M. J. Hopkins, D. F. Wright, T. Stadler, and R. C. Warnock. 2020b. Ignoring fossil age uncertainty leads to inaccurate topology and divergence time estimates in time calibrated tree inference. Frontiers in Ecology and Evolution 8:183.

Bogri, A., A. Solodovnikov, and D. Ż yla. 2018. Baltic amber impact on historical biogeography and palaeoclimate research: Oriental rove beetle Dysanabatium found in the Eocene of Europe (Coleoptera, Staphylinidae, Paederinae). Papers in Palaeontology 4:433–452.

Bouckaert, R., J. Heled, D. Kühnert, T. Vaughan, C.-H. Wu, D. Xie, M. A. Suchard, A. Rambaut, and A. J. Drummond. 2014. BEAST 2: A software platform for Bayesian evolutionary analysis. PLoS Computational Biology 10:e1003537.

Bouckaert, R., T. G. Vaughan, J. Barido-Sottani, S. Duchêne, M. Fourment, A. Gavryushkina, J. Heled, G. Jones, D. Kühnert, N. De Maio, et al. 2019. BEAST 2.5: An advanced software platform for Bayesian evolutionary analysis. PLoS Computational Biology 15:e1006650.

Brunke, A. J., S. Chatzimanolis, B. D. Metscher, K. Wolf-Schwenninger, and A. Solodovnikov. 2017. Dispersal of thermophilic beetles across the intercontinental Arctic forest belt during the early Eocene. Scientific Reports 7:12972.

Chen, W.-C. 2011. Overlapping Codon Model, Phylogenetic Clustering, and Alternative Partial Expectation Conditional Maximization Algorithm. Ph.D. thesis Iowa State University.

Cole, T. L., D. T. Ksepka, K. J. Mitchell, A. J. Tennyson, D. B. Thomas, H. Pan, G. Zhang, N. J. Rawlence, J. R. Wood, P. Bover, et al. 2019. Mitogenomes uncover extinct penguin taxa and reveal island formation as a key driver of speciation. Molecular Biology and Evolution 36:784–797.

Degrange, F. J., D. T. Ksepka, and C. P. Tambussi. 2018. Redescription of the oldest crown clade penguin: Cranial osteology, jaw myology, neuroanatomy, and phylogenetic affinities of madrynornis mirandus. Journal of Vertebrate Paleontology 38:e1445636.

Drummond, A. J., S. Y. Ho, M. J. Phillips, and A. Rambaut. 2006. Relaxed phylogenetics and dating with confidence. PLoS Biology 4:e88.

Drummond, A. J. and T. Stadler. 2016. Bayesian phylogenetic estimation of fossil ages. Philosophical Transactions of the Royal Society B: Biological Sciences 371:20150129.

Eguchi, S. and M. N. Tamura. 2016. Evolutionary timescale of monocots determined by the fossilized birth-death model using a large number of fossil records. Evolution 70:1136–1144.

Farrell, B. D. and A. S. Sequeira. 2004. Evolutionary rates in the adaptive radiation of beetles on plants. Evolution 58:1984–2001.

Gavryushkina, A., D. Welch, T. Stadler, and A. J. Drummond. 2014. Bayesian inference of sampled ancestor trees for epidemiology and fossil calibration. PLoS Computational Biology 10:e1003919.

Gradstein, F. M., J. G. Ogg, M. D. Schmitz, and G. M. Ogg. 2012. The Geologic Time Scale 2012. Elsevier.

Gustafson, G. T., A. A. Prokin, R. Bukontaite, J. Bergsten, and K. B. Miller. 2017. Tip-dated phylogeny of whirligig beetles reveals ancient lineage surviving on Madagascar. Scientific Reports 7:1–9.

Heath, T. A., J. P. Huelsenbeck, and T. Stadler. 2014. The fossilized birth-death process for coherent calibration of divergence-time estimates. Proceedings of the National Academy of Sciences of the United States of America 111:E2957–66.

Höhna, S., M. J. Landis, T. A. Heath, B. Boussau, N. Lartillot, B. R. Moore, J. P. Huelsenbeck, and F. Ronquist. 2016. RevBayes: Bayesian phylogenetic inference using graphical models and an interactive model-specification language. Systematic Biology 65:726–736.

Höhna, S., T. Stadler, F. Ronquist, and T. Britton. 2011. Inferring speciation and extinction rates under different sampling schemes. Molecular Biology and Evolution 28:2577–2589.

Holland, S. M. 2016. The non-uniformity of fossil preservation. Philosophical Transactions of the Royal Society B: Biological Sciences 371:20150130.

Jouault, C., F. Legendre, P. Grandcolas, and A. Nel. 2021. Revising dating estimates and the antiquity of eusociality in termites using the fossilized birth–death process. Systematic Entomology 46:592–610.

Kurumada, Y., S. Aoki, K. Aoki, D. Kato, M. Saneyoshi, K. Tsogtbaatar, B. F. Windley, and S. Ishigaki. 2020. Calcite U–Pb age of the Cretaceous vertebrate-bearing Bayn Shire Formation in the Eastern Gobi Desert of Mongolia: Usefulness of caliche for age determination. Terra Nova 32:246–252.

Lepage, T., D. Bryant, H. Philippe, and N. Lartillot. 2007. A general comparison of relaxed molecular clock models. Molecular Biology and Evolution 24:2669–2680.

Lewis, P. O. 2001. A likelihood approach to estimating phylogeny from discrete morphological character data. Systematic Biology 50:913–925.

Li, F., W. Ma, F. Meng, and H. Diao. 2021. Geochemical characteristics and geological significance of Daohugou Formation at Ningcheng County of Inner Mongolia, Eastern China. Geological Journal 56:2223–2239.

Pennell, M. W., J. M. Eastman, G. J. Slater, J. W. Brown, J. C. Uyeda, R. G. FitzJohn, M. E. Alfaro, and L. J. Harmon. 2014. geiger v2.0: an expanded suite of methods for fitting macroevolutionary models to phylogenetic trees. Bioinformatics 30:2216–2218.

Peppe, D. and A. Deino. 2013. Dating rocks and fossils using geologic methods. Nature Education Knowledge 4:1.

Pett, W. and T. A. Heath. 2020. Inferring the timescale of phylogenetic trees from fossil data. Pages 5.1:1–5.1:18 in Phylogenetics in the Genomic Era (C. Scornavacca, F. Delsuc, and N. Galtier, eds.). No commercial publisher | Authors open access book.

Pyron, R. A. 2011. Divergence time estimation using fossils as terminal taxa and the origins of Lissamphibia. Systematic Biology 60:466–481.

Rambaut, A., A. J. Drummond, D. Xie, G. Baele, and M. A. Suchard. 2018. Posterior summarization in Bayesian phylogenetics using Tracer 1.7. Systematic Biology 67:901–904.

Rambaut, A. and N. C. Grassly. 1997. Seq-Gen: an application for the Monte Carlo simulation of DNA sequence evolution along phylogenetic trees. Bioinformatics 13:235–238.

Robinson, D. and L. Foulds. 1981. Comparison of phylogenetic trees. Mathematical Biosciences 53:131–147.

Ronquist, F., S. Klopfstein, L. Vilhelmsen, S. Schulmeister, D. L. Murray, and A. P. Rasnitsyn. 2012a. A totalevidence approach to dating with fossils, applied to the early radiation of the Hymenoptera. Systematic Biology 61:973–999.

Ronquist, F., M. Teslenko, P. van der Mark, D. L. Ayres, A. Darling, S. Höhna, B. Larget, L. Liu, M. A. Suchard, and J. P. Huelsenbeck. 2012b. MrBayes 3.2: Efficient Bayesian phylogenetic inference and model choice across a large model space. Systematic Biology 61:539–542.

Schuster, A., S. Vargas, I. S. Knapp, S. A. Pomponi, R. J. Toonen, D. Erpenbeck, and G. Wörheide. 2018. Divergence times in demosponges (Porifera): first insights from new mitogenomes and the inclusion of fossils in a birth-death clock model. BMC Evolutionary Biology 18:1–11.

Shapiro, B., S. Y. Ho, A. J. Drummond, M. A. Suchard, O. G. Pybus, and A. Rambaut. 2011. A Bayesian phylogenetic method to estimate unknown sequence ages. Molecular Biology and Evolution 28:879–887.

Šmíd, J. and K. A. Tolley. 2019. Calibrating the tree of vipers under the fossilized birth-death model. Scientific Reports 9:1–10.

Stadler, T. 2010. Sampling-through-time in birth–death trees. Journal of Theoretical Biology 267:396–404.

Stadler, T. 2011. Simulating trees with a fixed number of extant species. Systematic Biology 60:676–684.

Stadler, T., A. Gavryushkina, R. C. Warnock, A. J. Drummond, and T. A. Heath. 2018. The fossilized birth-death model for the analysis of stratigraphic range data under different speciation modes. Journal of Theoretical Biology 447:41–55.

Stadler, T., D. Kühnert, S. Bonhoeffer, and A. J. Drummond. 2013. Birth–death skyline plot reveals temporal changes of epidemic spread in hiv and hepatitis c virus (hcv). Proceedings of the National Academy of Sciences 110:228–233.

Thomas, D. B., A. J. D. Tennyson, R. P. Scofield, T. A. Heath, W. Pett, and D. T. Ksepka. 2020. Ancient crested penguin constrains timing of recruitment into seabird hotspot. Proceedings of the Royal Society B: Biological Sciences 287:20201497.

Thorne, J., H. Kishino, and I. S. Painter. 1998. Estimating the rate of evolution of the rate of molecular evolution. Molecular Biology and Evolution 15:1647–1657.

Wang, X., Z. Zhou, H. He, F. Jin, Y. Wang, J. Zhang, Y. Wang, X. Xu, and F. Zhang. 2005. Stratigraphy and age of the daohugou bed in ningcheng, inner mongolia. Chinese Science Bulletin 50:2369–2376.

Zhang, C., T. Stadler, S. Klopfstein, T. A. Heath, and F. Ronquist. 2016. Total-evidence dating under the fossilized birth–death process. Systematic biology 65:228–249.

