## Supplementary methods and results for "Estimating the age of poorly dated fossil specimens and deposits using a total-evidence approach and the fossilized birth-death process"

### Supplementary materials

#### 1 Supplementary results on simulated data

Figure 1 shows the accuracy of extant tree reconstruction and tree height estimates on the simulated datasets. Based on these results, we can see that the extant tree topology is well estimated across all simulation conditions. The most-recent common ancestor (MRCA) age estimates for the full and extant trees are also accurate, although better for the full trees than the extant trees. Both MRCA age estimates are dependent on the simulation conditions, with accuracy decreasing when either the proportion of imprecise-date fossils or the relative age range of precise-date fossils increases.

#### 2 Alternative simulation conditions

In addition to the simulation datasets presented in the main text, we also studied the impact of several alternative simulation conditions:

1. all fossilization rates were divided by 2, which resulted in simulated trees containing between 20 and 30 fossils.
2. molecular sequences were simulated using a relaxed exponential clock with a mean rate of 0.005, identical to the rate used for the strict clock in the main datasets.
3. this condition replicated one common feature of insect morphological matrices, where some precise-date fossils are very incomplete and thus contain little data. In this condition a random sample of 5% of the precise-date fossils were assigned to be incomplete. A randomly chosen 10% of the characters in the matrix had data for these incomplete fossils, all other characters were assigned the unknown character state "?".
4. the morphological clock rate was set to 0.01, i.e. 10x lower than the main simulation.
5. no deposit was included, so all fossil specimens were treated as precise-date fossils.

6. the deposit was set to an interval of length 2 My, selected uniformly at random between 30 and 50 Ma. All imprecise-date fossils were thus sampled at approximately the same time.

7. the morphological clock model was set to a relaxed exponential clock with a mean clock rate of 0.1, identical to the rate used for the strict clock in the main simulation.

Alternative simulation conditions 1-3 were simulated with different proportions of imprecise-date fossils (0.1, 0.3 or 0.5) and using an age range of 0.2 times the true age for precise-date fossils. Alternative simulation conditions 4-7 were simulated with a 10% proportion of imprecise-date fossils (or 0% for the condition with no deposit) and using an age range of 0.1 times the true age for precise-date fossils. The inference was run identical to the main simulation, except for the clock models which were set to match the simulation clock models in conditions 2 and 7.

Results for conditions 1-3 are shown in Figures 2 and 3. Overall, these alternative simulation conditions did not affect the results much, with two notable exceptions. The coverage of the fossil age estimates was improved with lower numbers of fossils compared to the standard dataset, for both precise-date and imprecise-date fossils, although the relative error was similar to the standard. Applying a relaxed clock to the molecular alignment also appeared to decrease the accuracy of the topological placement of fossil samples compared to the standard dataset, although it should be noted that a high proportion of analyses using the relaxed clock and the 0.5 proportion of imprecise-date fossils failed to converge (34 out of 100 replicates).

Results for conditions 4-6 are shown in Figures 4 and 5. Results for condition 7 are not shown, as this analysis suffered from many convergence issues and not enough replicates converged for it to be included. Again, we see overall little effect of these alternative simulation conditions, although they slightly improve age estimates for precise-date fossils. The lower morphological clock rate also results in better topological estimates, as measured by the RF distance.

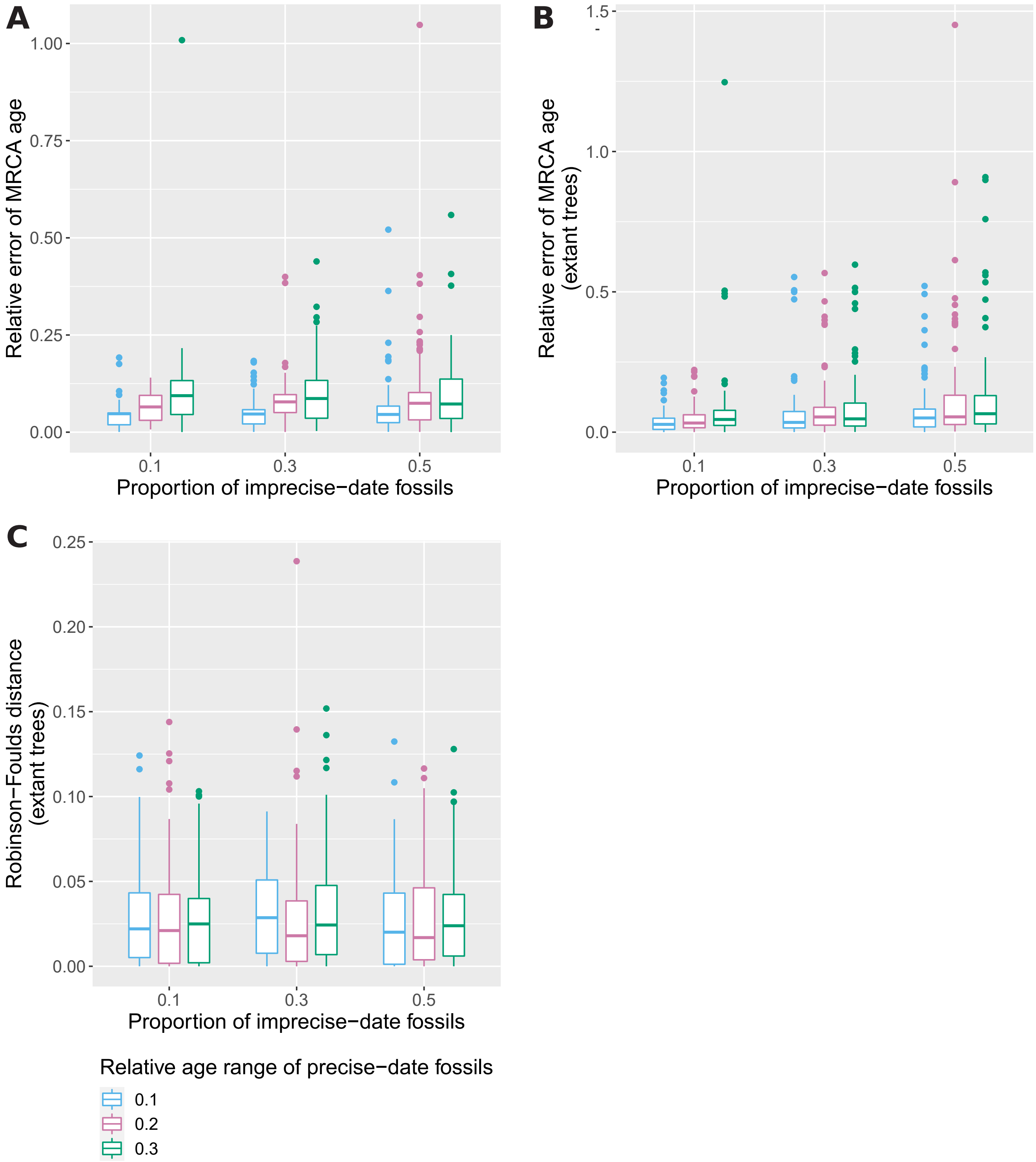


Figure 1: Mean normalized RF distance between estimated extant trees and simulated extant tree (A), relative error of the median MRCA age estimate (B) and relative error of the median MRCA age estimate for extant trees (C), for different proportions of imprecise-date fossils and different relative age ranges for precise-date fossils. Measures are averaged over all fossils for each replicate. The average and standard deviation across all replicates is shown.

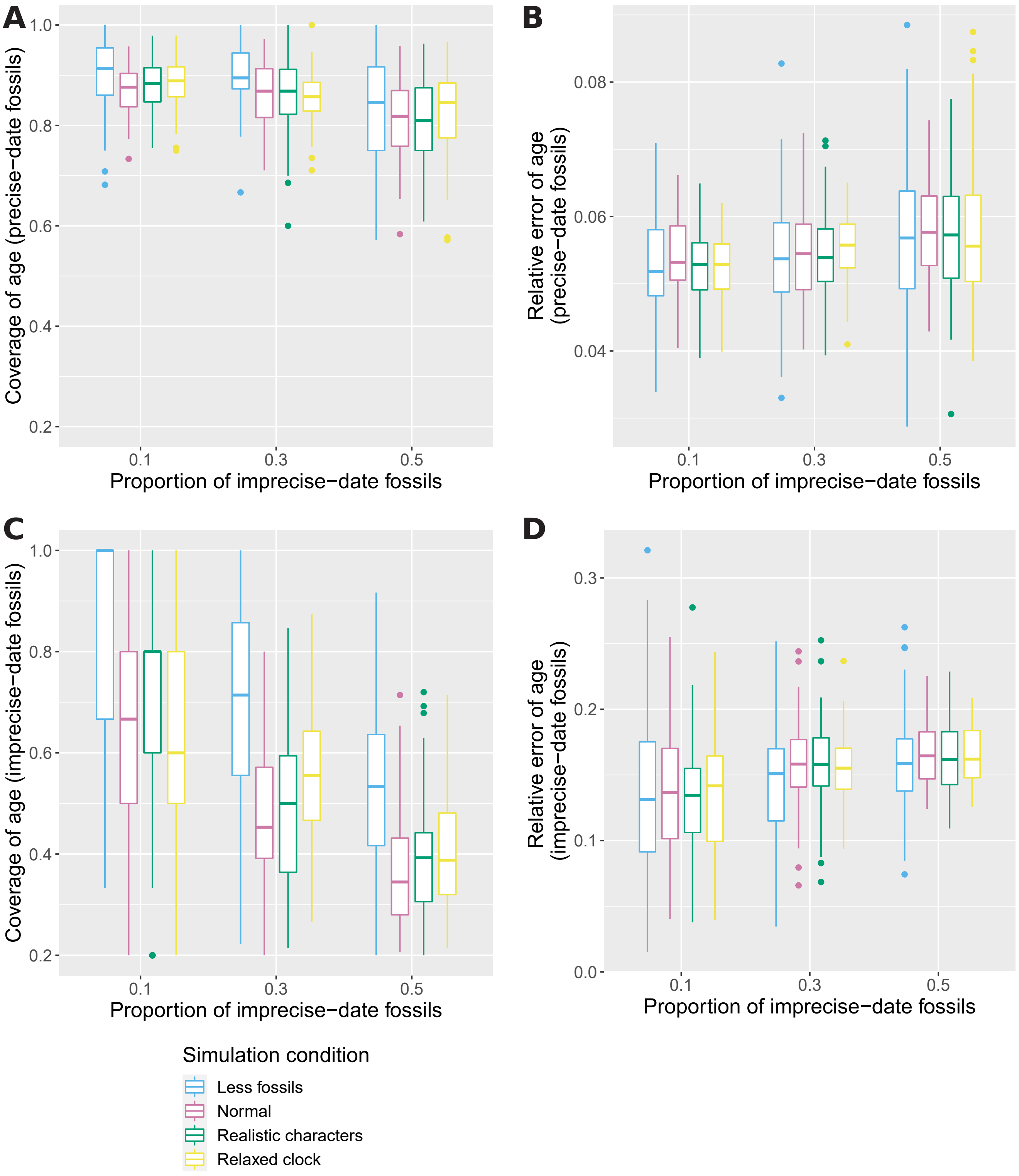


Figure 2: Relative error of the median age estimate (B,D) and 95% HPD coverage (A,C) of precise-date fossils (A,B) and imprecise-date fossils (C,D) for different proportions of imprecise-date fossils, and different simulation conditions. Measures are averaged over all fossils for each replicate. The average and standard deviation across all replicates is shown.

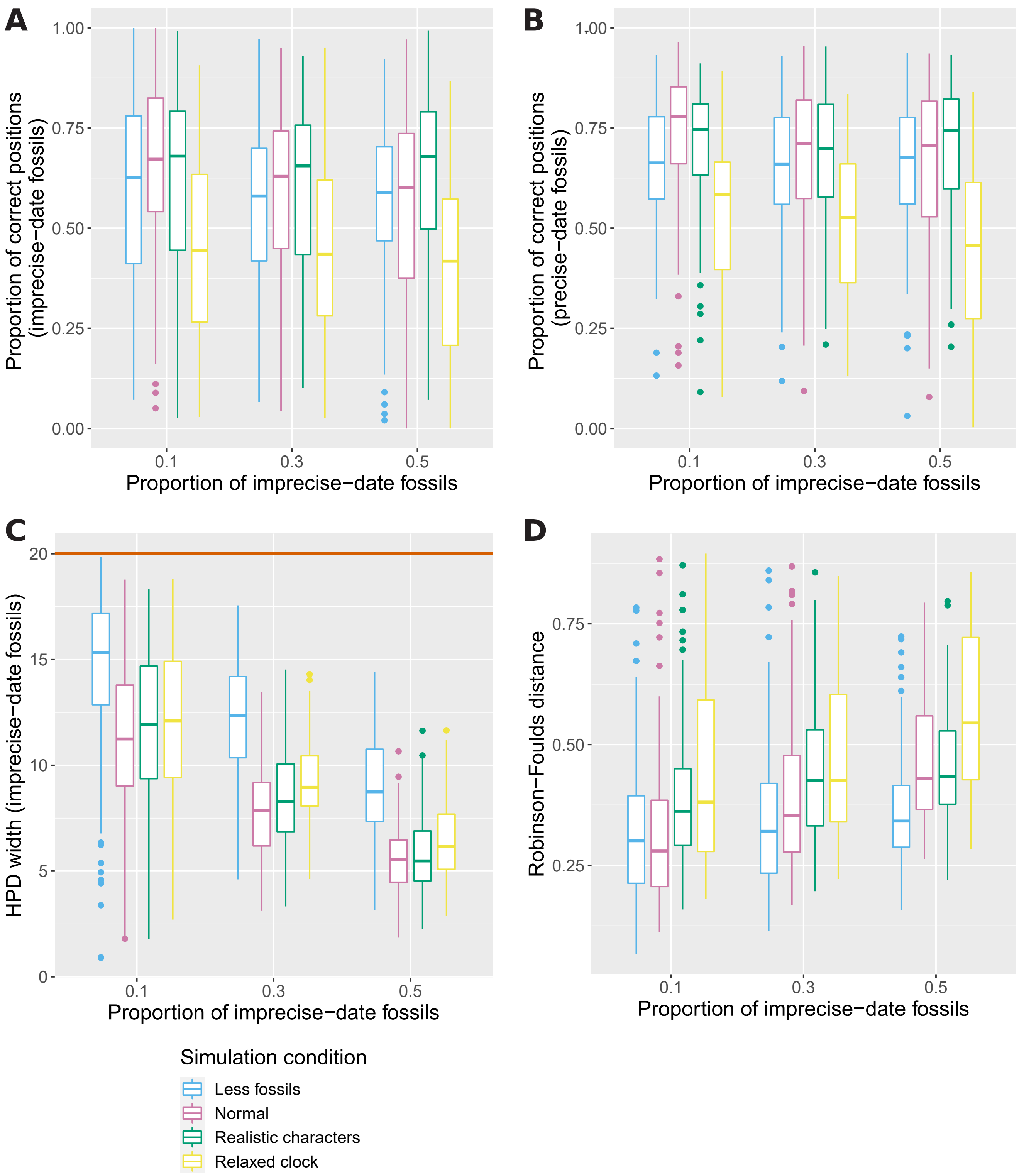


Figure 3: Proportion of posterior samples with correctly placed fossils, averaged across all precise-date fossils (A) or all imprecise-date fossils (B), width of the 95% HPD interval averaged across all imprecise-date fossils (C) and mean normalized RF distance between estimated trees and simulated tree (D), for different proportions of imprecise-date fossils and different simulation conditions. The average and standard deviation across all replicates is shown. The brown line in C shows the size of the age range set as prior for all imprecise-date fossils (*i.e.*, 20My).

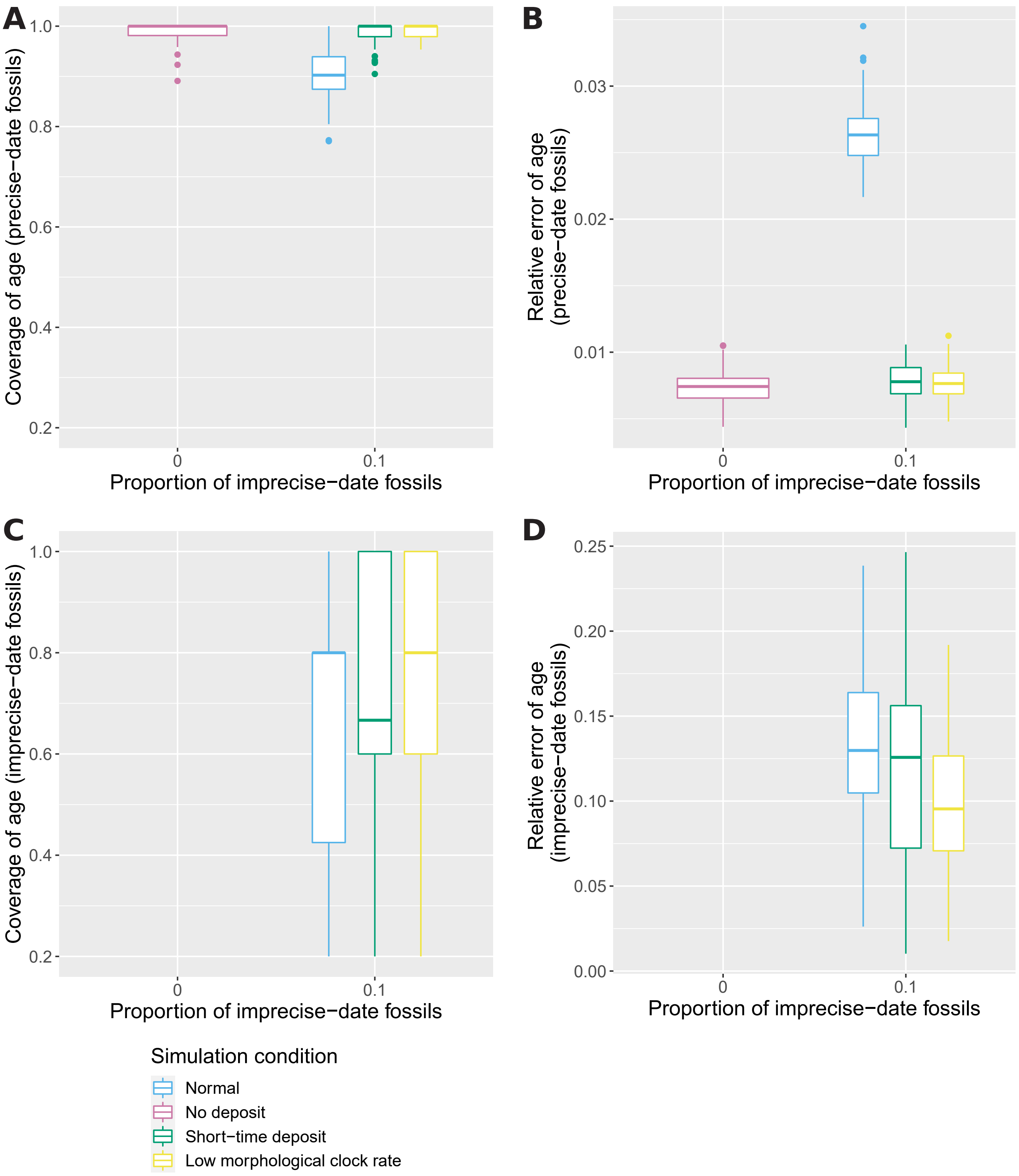


Figure 4: Relative error of the median age estimate (B,D) and 95% HPD coverage (A,C) of precise-date fossils (A,B) and imprecise-date fossils (C,D) for different simulation conditions. Measures are averaged over all fossils for each replicate. The average and standard deviation across all replicates is shown.

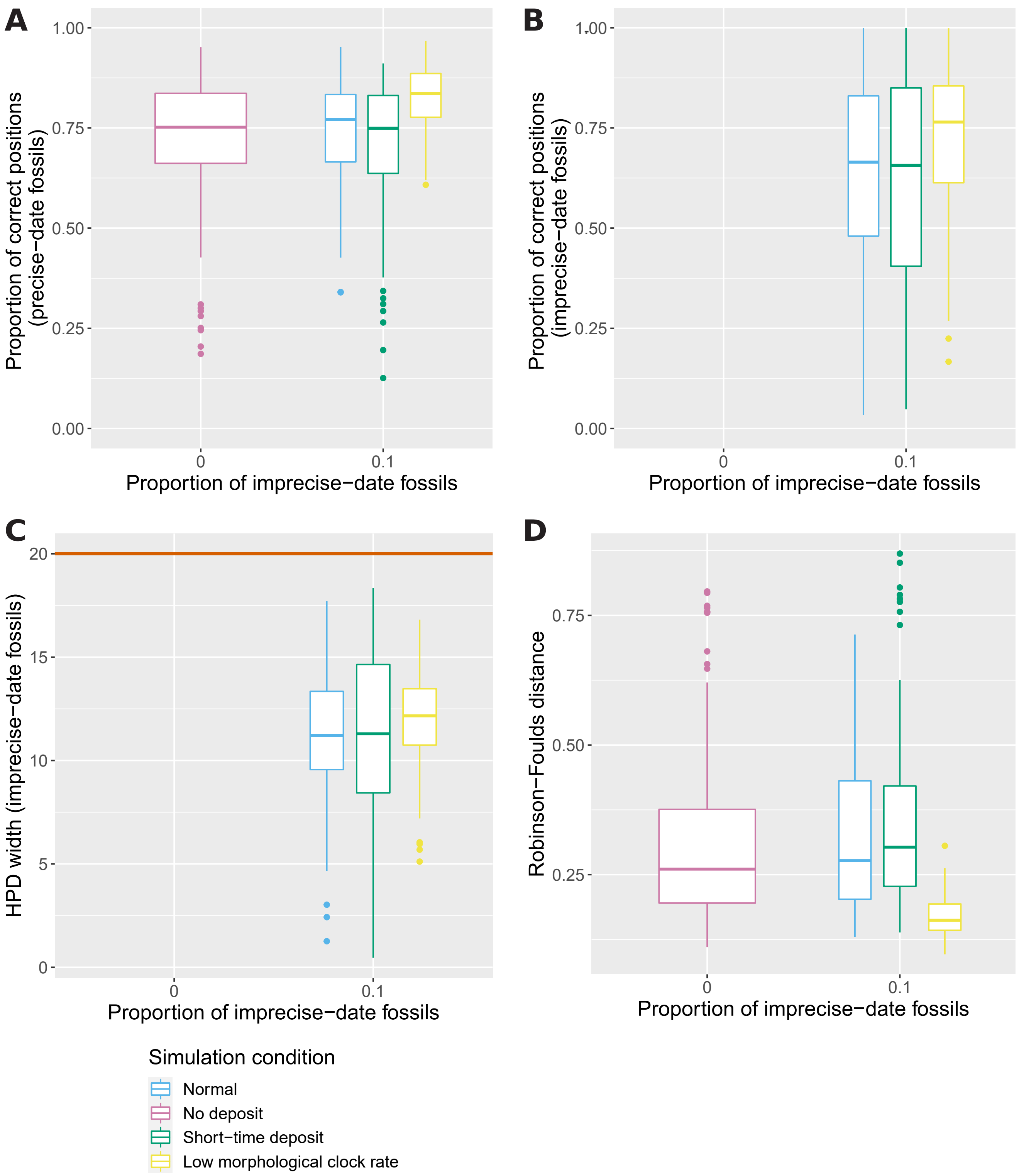


Figure 5: Proportion of posterior samples with correctly placed fossils, averaged across all precise-date fossils (A) or all imprecise-date fossils (B), width of the 95% HPD interval averaged across all imprecise-date fossils (C) and mean normalized RF distance between estimated trees and simulated tree (D), for different simulation conditions. The average and standard deviation across all replicates is shown. The brown line in C shows the size of the age range set as prior for all imprecise-date fossils (*i.e.*, 20My).

### 3 Empirical dataset

Figure 6 shows the results of the analysis on the penguins datasets, using either the small interval or its extension as deposit. Similar to the simulated datasets, increasing the number of imprecise-date fossils in the analysis decreases the accuracy of the estimates. In particular, when all the fossils in the extended interval are assigned to be imprecisely dated, the posterior distributions of ages for most fossils are extremely diffuse. The few posterior distributions which are well resolved are unreliable, as some (the *Kairuku* fossils) match well with the observed ranges, while others (the *Palaeeudyptes*) are completely different.

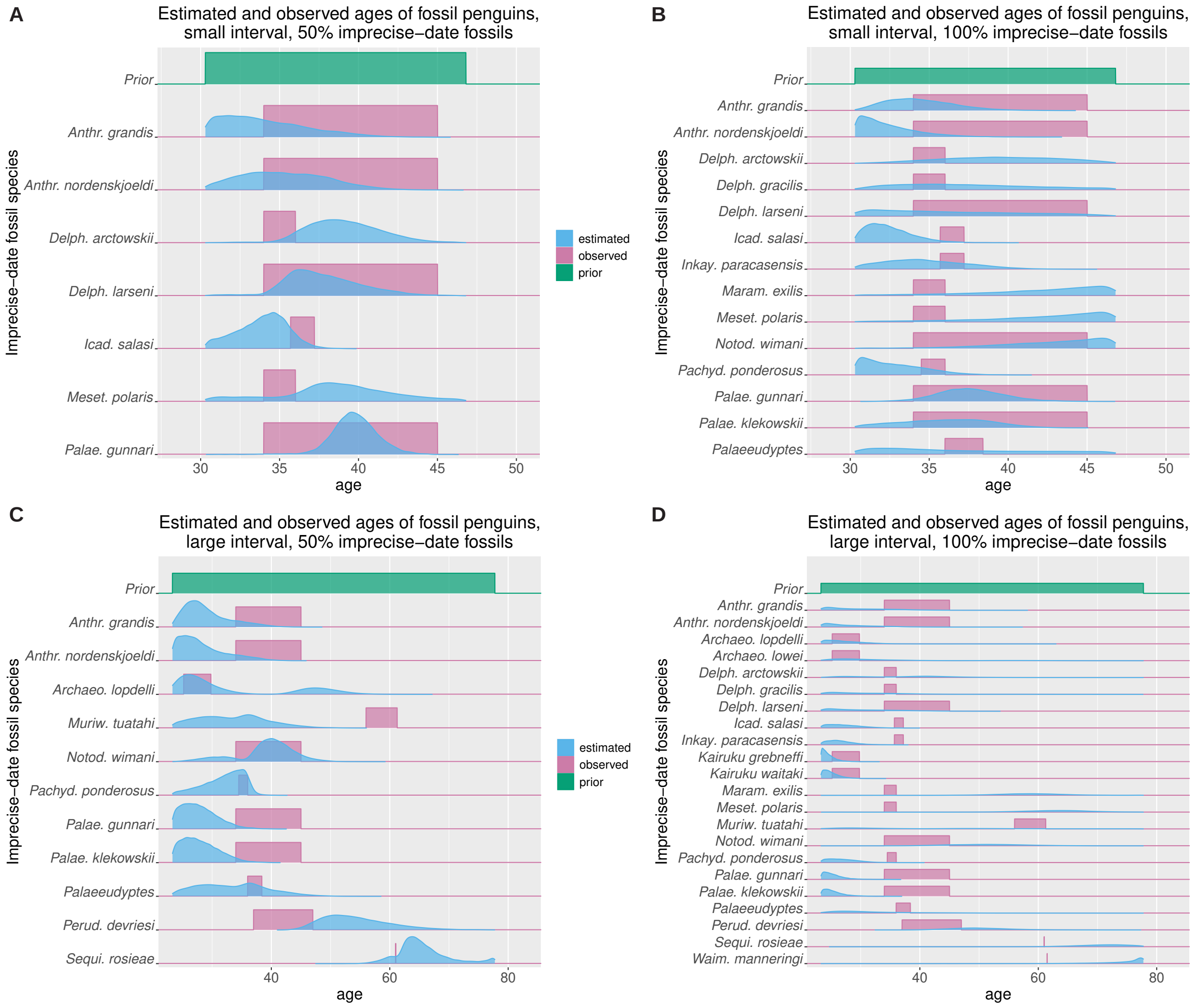


Figure 6: Comparison of observed (pink) and estimated (blue) penguin ages for the small (A,B) and extended (C,D) intervals, with a proportion of 0.5 (A,C) or 1 (B,D) of imprecise-date fossils. The observed age range is shown as a uniform distribution, while the estimated age is the inferred posterior distribution. The uniform distribution used as prior for the imprecise-date fossils is shown in green on each panel.

52 # 4 Supplementary figures

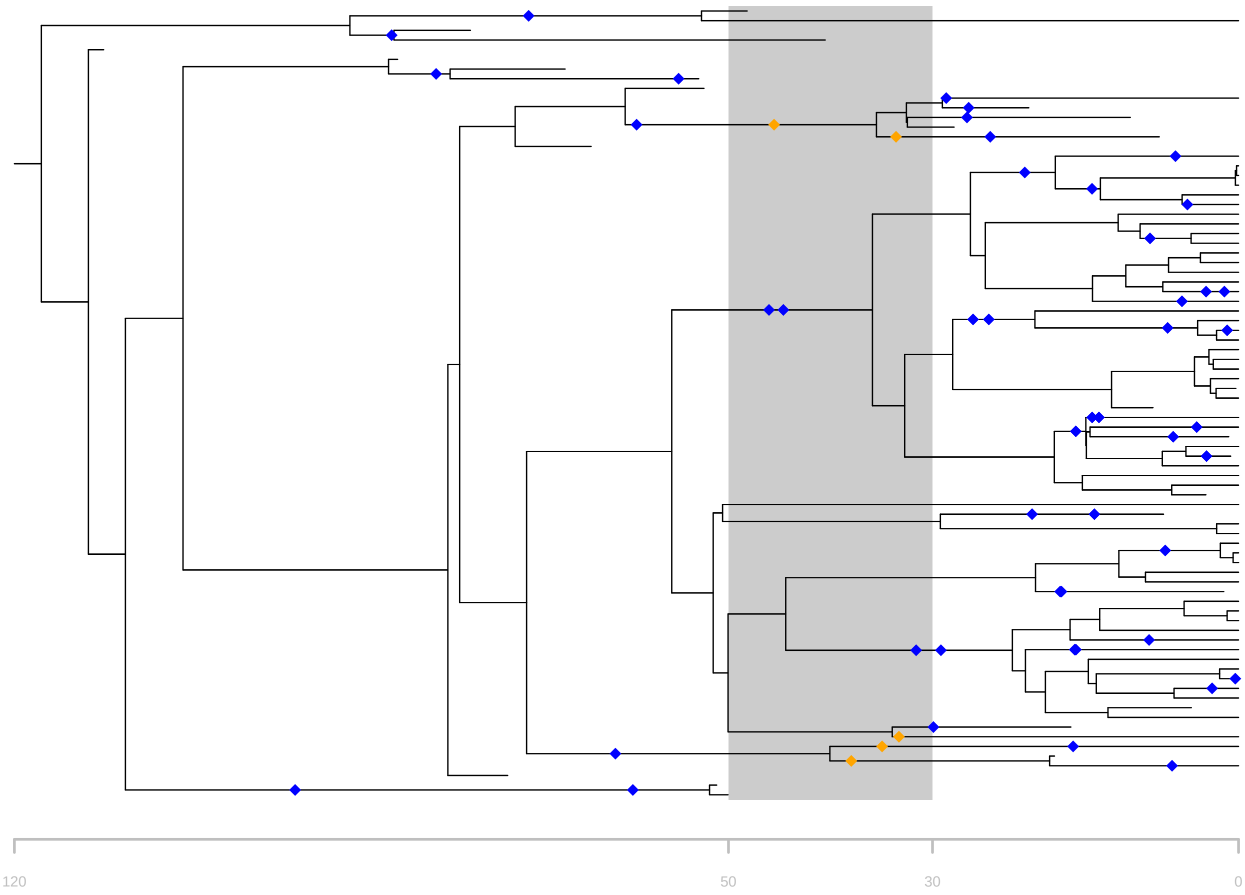


Figure 7: Example of a simulated (complete) tree with a 10% proportion of imprecise-date fossils. The undated deposit (from 30 to 50 Ma) is represented in dark gray. Fossil samples assigned to the imprecise-date deposit are shown in orange, fossils assigned to be precisely dated are shown in blue.

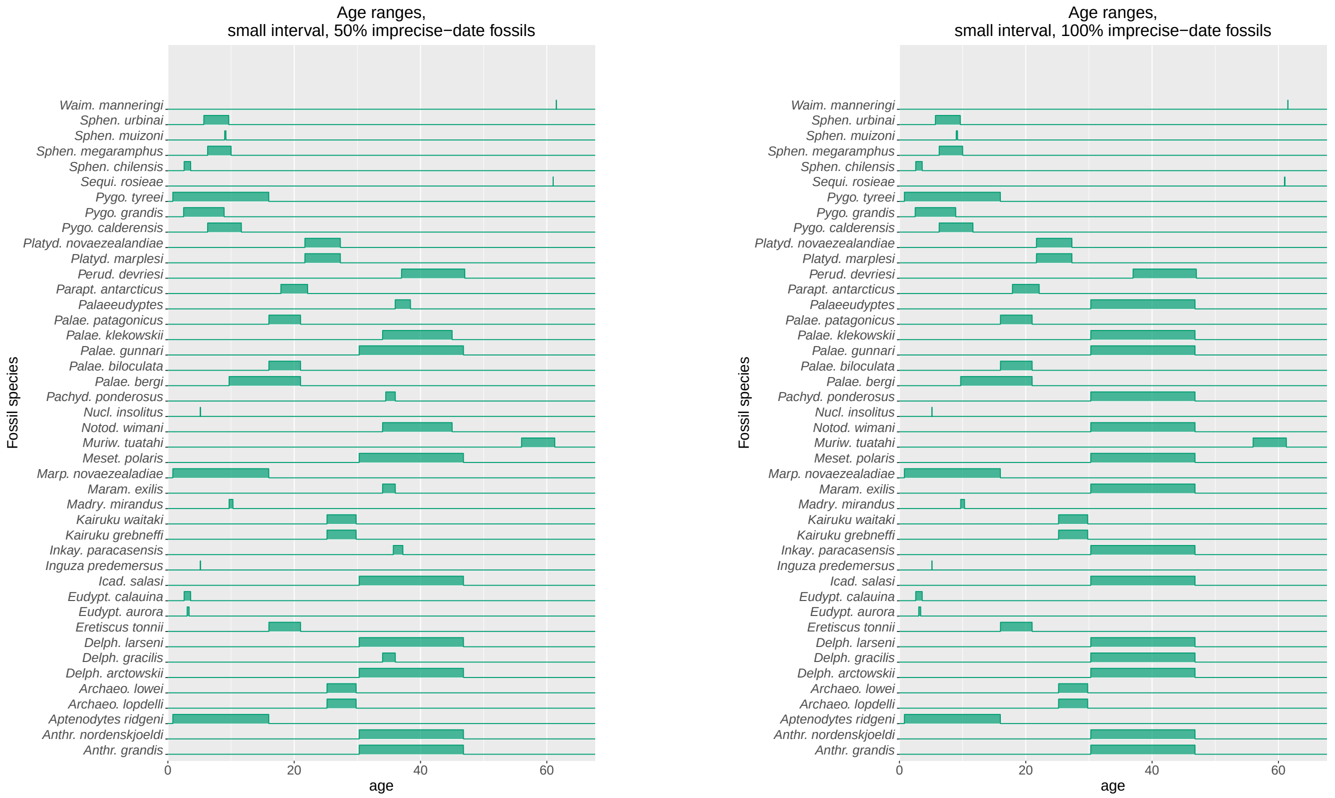


Figure 8: Prior age ranges set for each penguin fossil species using the small interval as imprecise-date deposit.

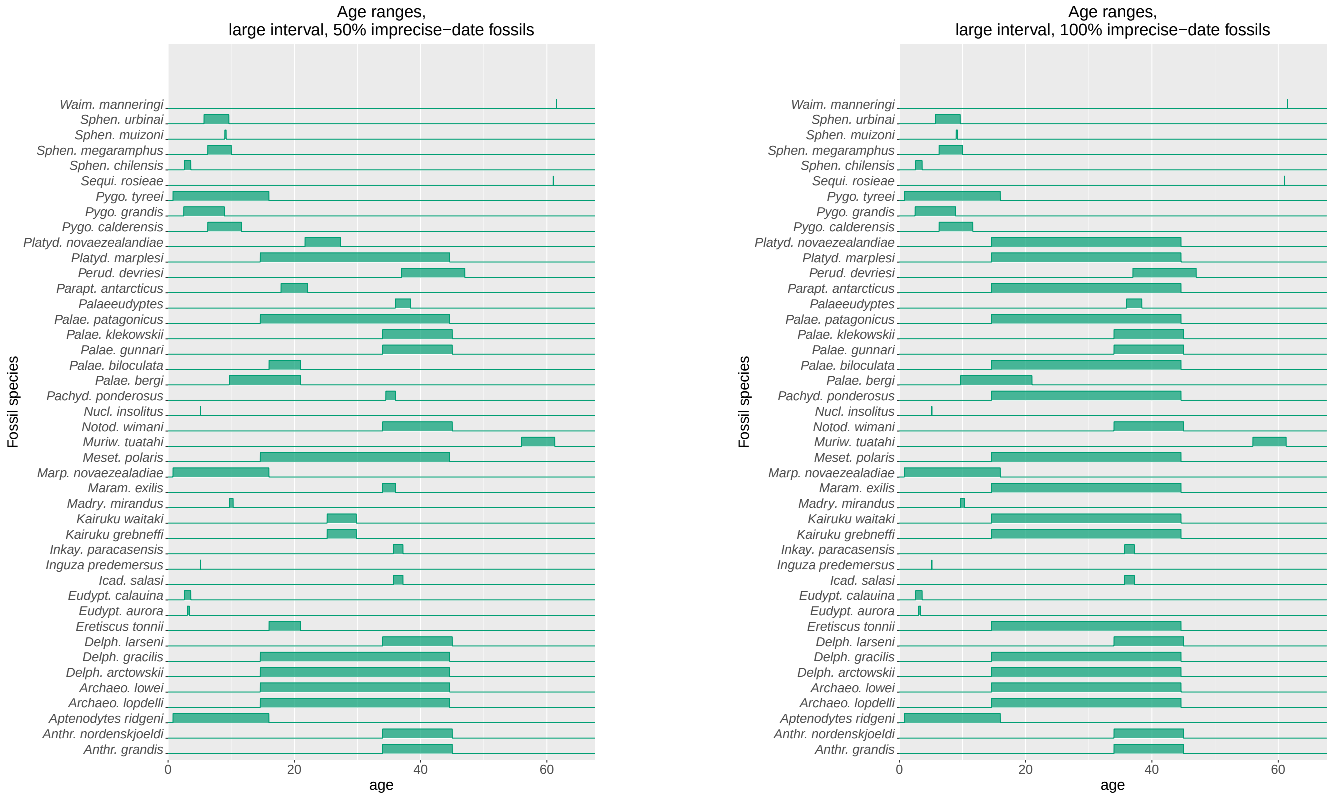


Figure 9: Prior age ranges set for each penguin fossil species using the large interval as imprecise-date deposit.

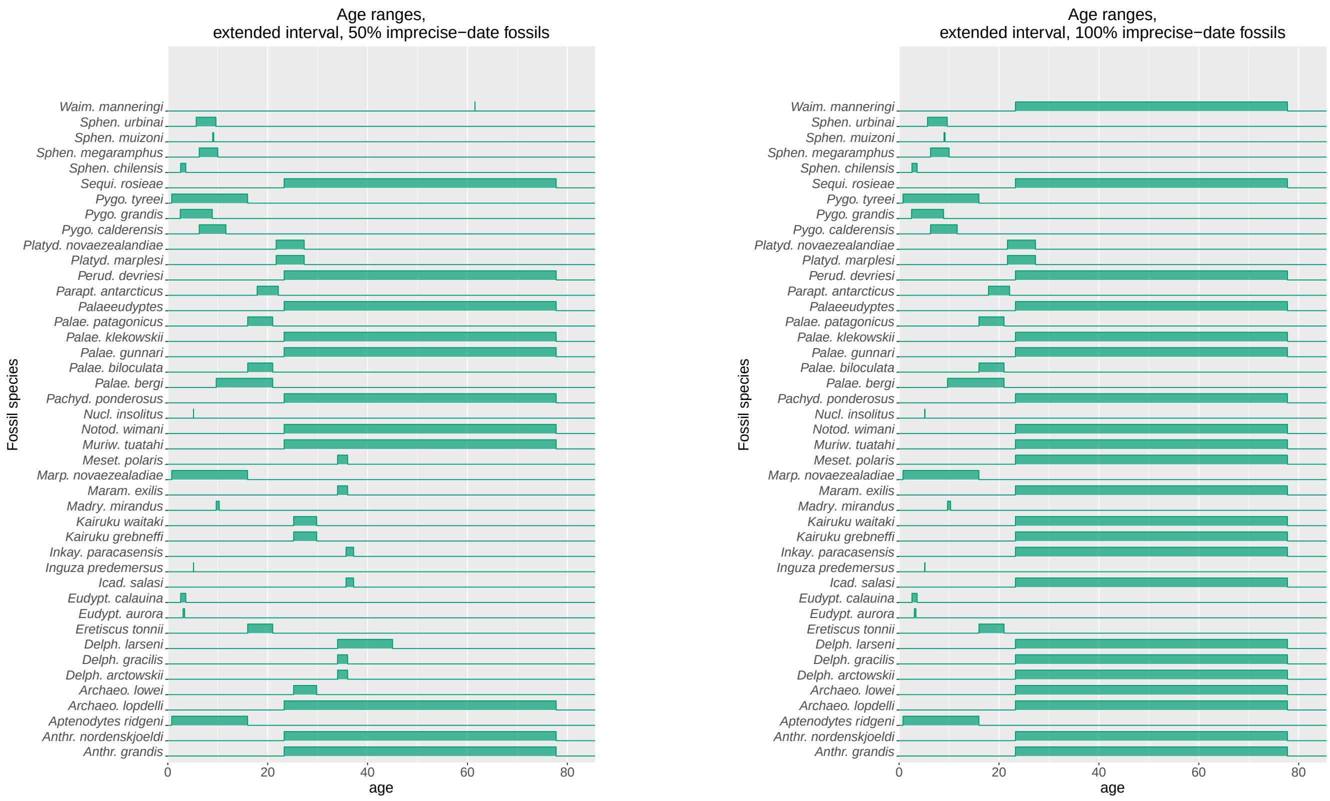


Figure 10: Prior age ranges set for each penguin fossil species using the extension of the small interval as imprecise-date deposit.

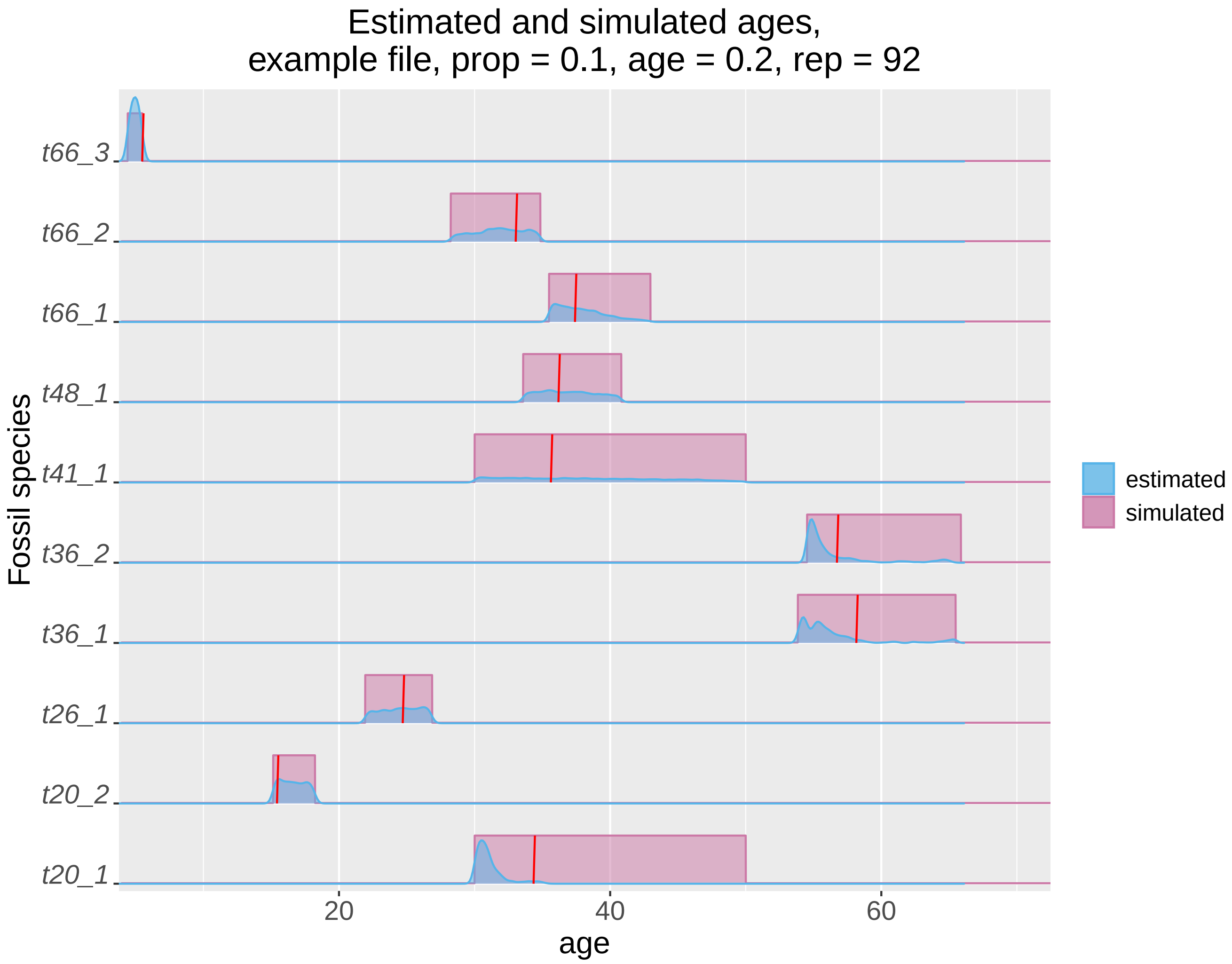


Figure 11: Comparison of simulated (pink) and estimated (blue) ages for one simulation replicate, with a 10% proportion of imprecise-date fossils and a relative age range of 0.2 for precise-date fossils. The simulated age range is shown as a uniform distribution, while the estimated age is the inferred posterior distribution. The true age of each fossil is marked in red.
